# Unsuspected involvement of spinal cord in Alzheimer Disease

**DOI:** 10.1101/673350

**Authors:** Roberta Maria Lorenzi, Fulvia Palesi, Gloria Castellazzi, Paolo Vitali, Nicoletta Anzalone, Sara Bernini, Elena Sinforiani, Giuseppe Micieli, Alfredo Costa, Egidio D’Angelo, Claudia A.M. Gandini Wheeler-Kingshott

## Abstract

**Objective:** Brain atrophy is an established biomarker for dementia, yet spinal cord involvement has not been investigated to date. As the spinal cord is relaying sensorimotor control signals from the cortex to the peripheral nervous system and viceversa, it is indeed a very interesting question to assess whether it is affected by atrophy in a disease that is known for its involvement of cognitive domains first and foremost, with motor symptoms being clinically assessed too. We therefore hypothesize that Alzheimer Disease severe atrophy can affect the spinal cord too and that spinal cord atrophy is indeed an important in vivo imaging biomarker contributing to understanding neurodegeneration associated with dementia.

**Methods:** 3DT1 images of 31 Alzheimer’s disease (AD) and 35 healthy control (HC) subjects were processed to calculate volumes of brain structures and cross-sectional area (CSA) and volume (CSV) of the cervical cord (per vertebra as well as the C2-C3 pair (CSA23 and CSV23)). Correlated features (ρ>0.7) were removed, and best subset identified for patients’ classification with the Random Forest algorithm. General linear model regression was used to find significant differences between groups (p<=0.05). Linear regression was implemented to assess the explained variance of the Mini Mental State Examination (MMSE) score as dependent variable with best features as predictors.

**Results:** Spinal cord features were significantly reduced in AD, independently of brain volumes. Patients classification reached 76% accuracy when including CSA23 together with volumes of hippocampi, left amygdala, white and grey matter, with 74% sensitivity and 78% specificity. CSA23 alone explained 13% of MMSE variance.

**Discussion:** Our findings reveal that C2-C3 spinal cord atrophy contributes to discriminate AD from HC, together with more established features. Results show that CSA23, calculated form the same 3DT1 scan as all other brain volumes (including right and left hippocampi), has a considerable weight in classification tasks warranting further investigations. Together with recent studies revealing that AD atrophy is spread beyond the temporal lobes, our result adds the spinal cord to a number of unsuspected regions involved in the disease. Interestingly, spinal cord atrophy explains also cognitive scores, which could significantly impact how we model sensorimotor control in degenerative diseases with a primary cognitive domain involvement. Prospective studies should be purposely designed to understand the mechanisms of atrophy and the role of the spinal cord in AD.

## 1 Introduction

Dementia is one of the most debilitating cognitive neurodegenerative disorders affecting the central nervous system in elderly people and having a significant impact on daily life activities. With an ageing population the incidence of dementia is growing and the consequences on society are huge. Clinically, several forms of dementia-like diseases that differently impair multiple cognitive and behavioral domains are defined. Alzheimer’s disease (AD) is the most common cause of dementia and it is responsible for 60% to 80 % of cases worldwide^1^. What is the effect of neurodegeneration on sensorimotor control is an interesting question because it is believed to be highly relevant also for understanding cognitive functions. As the spinal cord is relaying sensorimotor control signals from the cortex to the peripheral nervous system and viceversa, it is indeed important to assess whether it is affected by atrophy in a disease that is known for its involvement of cognitive domains. Recent indications suggest that there is definitely a sensorimotor network rewiring and that the motor system may even be affected before cognitive functions in AD^2–6^. Clinical symptoms of early AD include amongst others fine motor impairment, with for example worsening of writing abilities. Therefore, it is important to understand first of all whether the spinal cord plays a part in this disease and to understand how significant is its involvement.

AD is associated with an extracellular deposit of β-amyloid plaques in the brain and cerebral vessels, but also to the presence of intracellular neurofibrillary tangles, which appear like paired helical filaments with hyperphosphorylated tau proteins. Tau tangles have been identified as the cause of cortical neurons’ degeneration while β-amyloid oligomers have an important role in synaptic impairment, hence β-amyloid plaques deposition is suggested to raise later during the AD progression^7,8^. This neuronal degeneration explained by pathophysiology leads to macroscopic atrophy of specific brain structures, such as the hippocampi and the medial temporal lobes ^9^, which can be detected using Magnetic Resonance Imaging (MRI) techniques. Indeed, several MRI studies have demonstrated significant atrophy of white matter, gray matter and specific brain structures such as the hippocampi, thalami and amygdalae in AD patients suggesting that these structures are informative in identifying dementia disorders^10,11^. The hippocampi have been proposed as in vivo non-invasive imaging biomarkers of AD while other structures may be useful in distinguishing between different subtypes of dementia^12^. Only far and few old studies have looked at the spinal cord in AD, from a postmortem histochemical analysis and with reference to the autonomic system, but results were never reproduced or follow through as they focused on tau pathology, which was only sporadically reported^13^.

Recently, numerous MRI investigations have tried to identify new in vivo biomarkers for dementia to understand mechanisms of AD, to have better tools for assessing new therapies and predicting the clinical evolution of prodromic stages of dementia. Optical Coherence Tomography studies, for example, have been used to demonstrate that retinal ganglion cell degeneration can be associated to early stages of AD. Also, structures like the cerebellum, not classically associated with AD, have been found to be altered in imaging studies of dementia^2^, with atrophy of the anterior cerebellum - known for its motor control - being present even in the prodromic stages of mild cognitive impairment (MCI)^14^. A recent work has also looked at graph theory metrics to distinguish patterns of AD, identifying potentially different subtypes^15^, although focusing on cortical and deep grey matter areas, without including the cerebellum and the spinal cord. Studies of other diseases associated with neurodegeneration, such as multiple sclerosis^16^, amyotrophic lateral sclerosis^17^, and spinal cord injury^18^, have revealed that atrophy of the spinal cord is indicative of widespread alterations of the central nervous system and might be considered as a relevant imaging biomarker in a wider range of neurodegenerative diseases. Nevertheless, this kind of alteration has never been investigated and reported in dementia patients. Hence, the main aim of the present work was in the first instance to assess whether spinal cord volume is reduced in AD patients compared to healthy controls (HC), hypothesizing that the neurodegeneration typical of AD spreads to all components of the central nervous system; we achieved this by comparing a number of spinal cord features between AD and HC. This information is very important for our understanding of how a neurodegenerative disease like AD has implications beyond the known brain atrophy: this could also have significant impact on future modeling of brain networks. Furthermore, in case of a positive outcome, it is important to quantify the role of spinal cord features in distinguishing between AD and HC to drive the design of future studies; for this we implemented a machine learning approach for features selection, that is increasingly applied to improve diagnostic accuracy by quantitative imaging^19,20^. Finally, we quantified the contribution of spinal cord atrophy to explain variance of clinical scores for determining its clinical relevance.

## 2 Materials and Methods

### 2.1 Subjects

A total of 66 subjects including 31 AD patients (age (73 ± 7) years, 12 females (F), MMSE = 16 ± 6) and 35 HC (age (69 ± 10) years, 17 F, MMSE = 28 ± 1), as a reference group, were analyzed. 7 subjects (4 HC and 3 AD) were excluded from the study due to post-processing issues, hence the final dataset comprised 32 HC and 28 AD.

Inclusion criteria for patients were: clinical diagnosis of dementia on the basis of the Diagnostic and Statistical Manual of Mental Disorders (DSM-5) criteria^21^, Mini-Mental State Examination (MMSE) score^22^ below 24 and age above 60 years. Exclusion criteria comprised the presence of at least one of the following: epilepsy or isolated seizures, major psychiatric disorders over the previous 12 months, pharmacologically treated delirium or hallucinations, ongoing alcoholic abuse, acute ischemic or hemorrhagic stroke, known intracranial lesions, and systemic causes of subacute cognitive impairment^23^. Diagnosis of AD was made according to the criteria of the National Institute of Neurological and Communicative Disorders and Stroke and Alzheimer’s Disease and Related Disorders Association (NINCDS-ADRDA) workgroup^24^. HC were enrolled on a voluntary basis among subjects with MMSE score above 27 and attending a local third age university (University of Pavia, Information Technology course) or included in a program on healthy ageing (Fondazione Golgi, Abbiategrasso, Italy).

The study was accomplished in accordance with the Declaration of Helsinki and with the approbation of the local ethic committee of the IRCCS Mondino Foundation, upon signature of the written informed consent by the subjects.

### 2.2 MRI Acquisition

High resolution 3D T1-weighted(3DT1-w) MR images were acquired using a Siemens MAGNETOM Skyra3T(Siemens AG, Erlangen, Germany) with software version NUMARIS/4(syngo MR D13C version) and a receiving head-coil with 32 channels.

Scan parameters were^12^: TR=2300ms, TE=2.95ms, TI=900 ms, flip angle=9degrees, field of view (FOV)=269×252mm, acquisition matrix=256×240, in-plane resolution=1.05×1.05mm, slice thickness=1.2 mm, and 176 sagittal slices. The FOV, in feet-to-head direction, was set to cover the entire brain and cervical cord up to the C5 vertebra in all subjects.

### 2.3 Spinal Cord analysis

For each subject, the 3DT1-w volume (the same used normally for brain atrophy measurements - see below) was resized removing the brain and centering the FOV on the spine. Once a single volume of interest (VOI) comprising the same spinal cord regions for each 3DT1-w was defined (matrix=176×240×96 voxels), the process was automatized for the whole dataset. The resized 3DT1-w volumes were analyzed with the Spinal Cord Toolbox (http://sourceforge.net/projects/spinalcordtoolbox), an open source software specifically developed to elaborate spinal cord images, to extract features of the C1-C5 vertebrae.

The spinal cord was segmented with the *propseg* algorithm^25^ and manually labelled^26^ to identify all vertebrae separately^27^(Figure 1). Mean cross-sectional area (CSA) and volume (CSV) were calculated for each vertebra and for the C2-C3 pair^28,29^(CSA23 and CSV23), given the known sensitivity of this combined level to disease severity^30^. CSA is computed by counting pixels in each slice and then geometrically adjusting it multiplying by the angle (in degrees) between the spinal cord centerline and the inferior-superior direction. CSV, indeed, is computed by counting pixels and multiplying by slice thickness.

**Figure 1.**
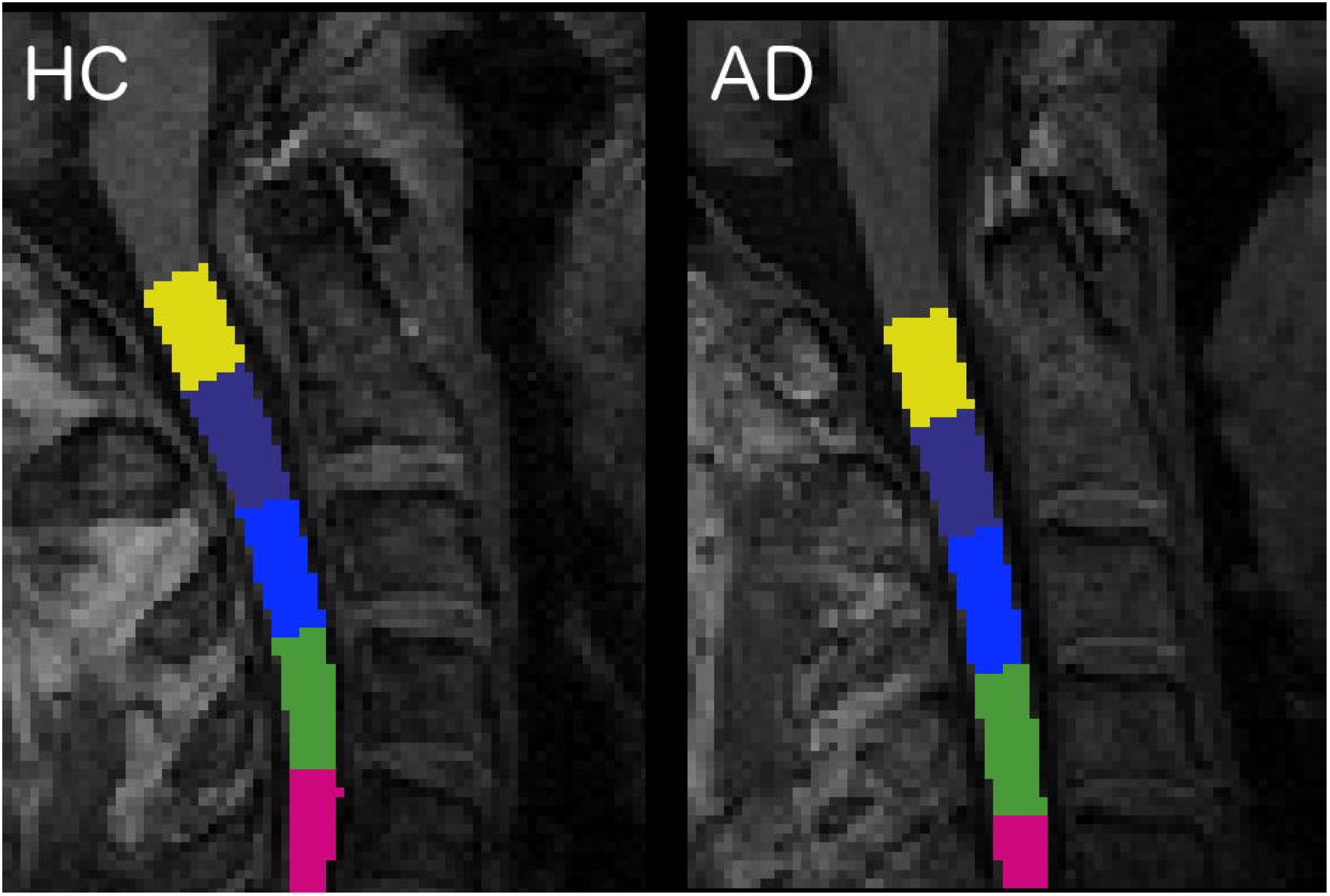
Labelled vertebrae in two randomly chosen subjects: a HC subject on the left and an AD patient on the right (slice n=96, sagittal plane). Each color represents a different vertebra from C1 (yellow) to C5 (fuchsia).

### 2.4 Brain atrophy analysis

The 3DT1-w images were also segmented into white matter (WM), gray matter (GM) and cerebrospinal fluid (CSF) using SPM12 (https://www.fil.ion.ucl.ac.uk/spm/software/spm12)^31^, while left (L) and right (R) hippocampi (LHip and RHip), thalami (LThal and RThal) and amygdalae (LAmy and RAmy) were segmented using FIRST (FSL, https://fsl.fmrib.ox.ac.uk/fsl/fslwiki/FIRST)^32^ (Figure 2).

**Figure 2.**
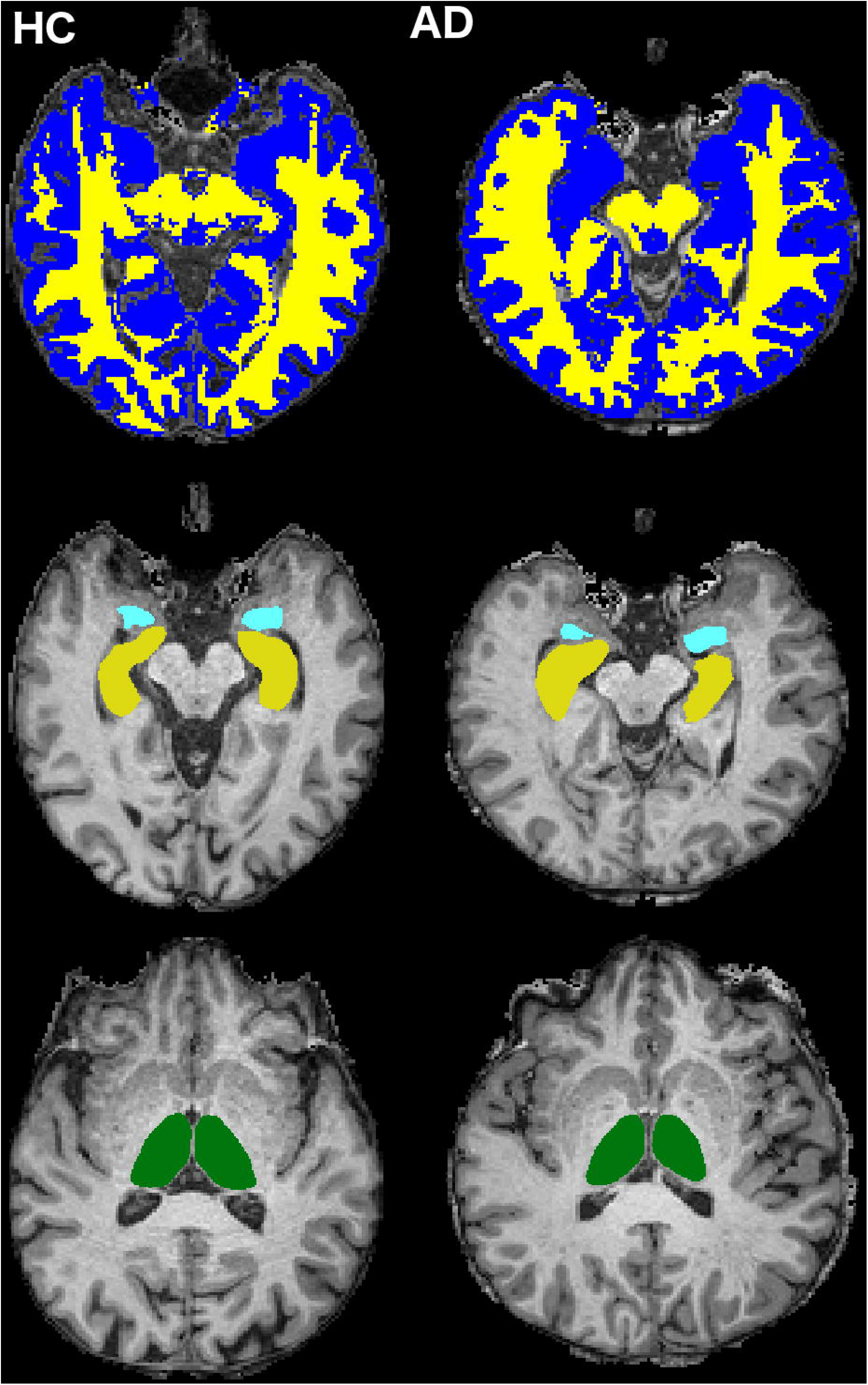
Cerebral tissue segmentation in two randomly chosen subjects: a HC subject on the left and an AD patient on the right. Top row: WM (yellow) and GM (blue) segmentation (slice n = 126, transverse plane). Middle row: hippocampi (yellow) and amygdalae (light blue) segmentation (slice n = 123, transverse plane). Bottom row: thalami (green) segmentation (slice n = 132, transverse plane).

WM, GM and all other brain structures volumes were calculated in mm^3^. Total intracranial volume, as the sum of WM, GM and CSF, was also calculated to account for different brain sizes.

### 2.5 Classification of AD and feature selection analysis

Classification between AD and HC was performed using a machine learning approach implemented in Orange (https://orange.biolab.si/).

A total of 22 features were extracted from the above MRI morphometric analysis. Given the large number of parameters extracted compared to the sample size of our AD and HC groups, a feature reduction approach was adopted in order to control for overfitting issues. The Spearman correlation coefficient^33^ was obtained in Matlab between pairs of all calculated metrics. When pairs of metrics had a correlation coefficient greater than 0.7, one metric was kept while the other was eliminated.

Ranking was implemented with the ReliefF algorithm^34^ on the uncorrelated features to identified the best subset able to classify AD from HC, and particularly to investigate the contribution of spinal cord metrics to the task. In order to identify a unique subset of features, 30% of instances was employed for ranking. Data were normalized by span to avoid a polarization of the results due to the different scale of features, as for WM compared to CSA. The remaining 70% of instances was further divided into 70% for the Random Forest algorithm application and 30% to test its classification accuracy (CA=(True Positive + True Negative)/(True Positive + True Negative + False Positive + False Negative)), sensitivity (Sens= True Positive/(True Positive + False negative)) and specificity (Spec= True Negative/(True Negative + False Positive)), using the previously-identified best features.

Among several machine learning algorithms, RF was selected for its robustness against a reduced number of input features and the capacity to weight features runtime, providing features relevance in a classification task^35,36^. The Receiving Operating Characteristics (ROC) curve was then obtained to visually discriminate between AD and HC and the Area Under the Curve was also calculated to quantify the overall ability of RF to discriminate between AD and HC.

### 2.6 Statistical analysis

Statistical tests were performed using the Statistical Package For Social Sciences (SPSS) software, version 21 (IBM, Armonk, New York). All continuous data were tested for normality using a Shapiro-Wilk test^37^. Age and MMSE were compared between AD and HC using a two-tailed Kruskal-Wallis test^38^ while gender was compared using a chi-squared test^39^. A multivariate regression model with gender, age and total intracranial volume as covariates was used to compare all morphometric metrics between AD and HC. Two-sided *p*<0.05 was considered statistically significant.

Furthermore, to assess the power of the best features in explaining the variance of the MMSE, a linear regression model was implemented using the MMSE score as the dependent variable and the best features as predictors. These independent features were used in two ways: i) each predictor was used alone to determine its specific contribution to MMSE; ii) all features were used in a backward approach to identify which of them explained the greatest percentage of MMSE variance. A threshold of *p*<0.01(two-tailed) was considered statistically significant.

## 3 Results

### 3.1 Subjects

Population demographics and neuropsychological scores are reported in Table 1. Significant differences were found in MMSE between HC and AD patients.

**Table 1:** Subjects’ demographic and neuropsychological data.

|  | HC (n=32) | AD (n=28) | p-value |
| --- | --- | --- | --- |
|  | mean (SD) | mean (SD) |  |
| Age [yrs] | 69.4 (9.6) | 73.0 (6.4) | 0.138 |
| Gender (Male [%]) | 51.4 | 56.2 | 0.800 |
| MMSE | 28.5 (0.2) | 16.0 (1.1) | < 0.001* |

Gender is expressed in Male % and compared with a Chi-square test. Age and MMSE are expressed as mean (SD) and compared with a Kruskal-Wallis test. Significance was set to p=0.05. \* refers to statistically significant comparisons.

### 3.2 Morphometric changes in AD patients

All results are reported in Table 2 and Table 3. AD patients compared to HC showed atrophy in all brain structures. Moreover, all patients for all investigated spinal cord segments showed reduced CSA at all vertebral levels, while CSV was significantly reduced only in correspondence of vertebrae C1 and C2.

**Table 2:** Brain morphometric changes in AD patients.

|  |  | HC (n=32) | AD (n=28) | p-value |
| --- | --- | --- | --- | --- |
|  |  | mean (SD) | mean (SD) |  |
| Brain Structures (mm<sup>2</sup>) | ICV | 1573086 (144439) | 1511611 (139532) | 0.04* |
|  | WM | 612335 (11230) | 540237 (12064) | <0.001* |
|  | GM | 427508 (6492) | 399274 (6975) | 0.006* |
|  | RHip | 3602 (106) | 2932 (114) | <0.001* |
|  | LHip | 3591 (99) | 2822 (107) | <0.001* |
|  | LThal | 7013 (109) | 6433 (118) | 0.001* |
|  | RThal | 6808 (109) | 6371 (117) | 0.011* |
|  | LAmy | 1256 (41) | 1054 (44) | 0.002* |
|  | RAmy | 1323 (63) | 1120 (66) | 0.035* |

Volumes of different brain structures expressed in mm<sup>3</sup>. Values are expressed as mean (SD). Significance was set at p = 0.050. \* refers to statistically significant values.

**Table 3:** Spinal cord morphometric changes in AD patients.

| Vertebra |  | HC (n=32) | AD (n=28) | p-value |
| --- | --- | --- | --- | --- |
|  |  | mean (SD) | mean (SD) |  |
| Area (mm<sup>2</sup>) | C1 | 69.8 (1.6) | 63.1 (1.8) | 0.009* |
|  | C2 | 65.7 (1.3) | 60.2 (1.4) | 0.008* |
|  | C3 | 62.5(1.4) | 56.9 (1.6) | 0.013* |
|  | C4 | 62.5 (1.6) | 57.2 (1.7) | 0.031* |
|  | C5 | 58.9 (1.6) | 52.8 (1.7) | 0.019* |
|  | C2-C3 | 65.1 (1.6) | 58.3 (1.7) | 0.007* |
| Volume (mm<sup>3</sup>) | C1 | 883.4 (27.3) | 800.4 (29.3) | 0.050* |
|  | C2 | 979.8 (28.4) | 857.1 (30.6) | 0.006* |
|  | C3 | 932.3 (29) | 886.9 (31.2) | 0.308 |
|  | C4 | 882.3 (35.1) | 807.9 (37.7) | 0.168 |
|  | C5 | 667 (34.5) | 609.1 (37.1) | 0.275 |
|  | C2-C3 | 1860.5 (66.8) | 1729.9 (71.7) | 0.204 |

Cross sectional area (in mm<sup>2</sup>) and volumes (in mm<sup>3</sup>) of spinal cord vertebrae. Values are expressed as mean (SD). Significance was set at p = 0.050. \* refers to statistically significant values.

### 3.3 AD classification based on morphometric data

Results of the correlation analysis are reported in Figure 3, and show that brain volumes are not significantly correlated with spinal cord metrics.

**Figure 3.**
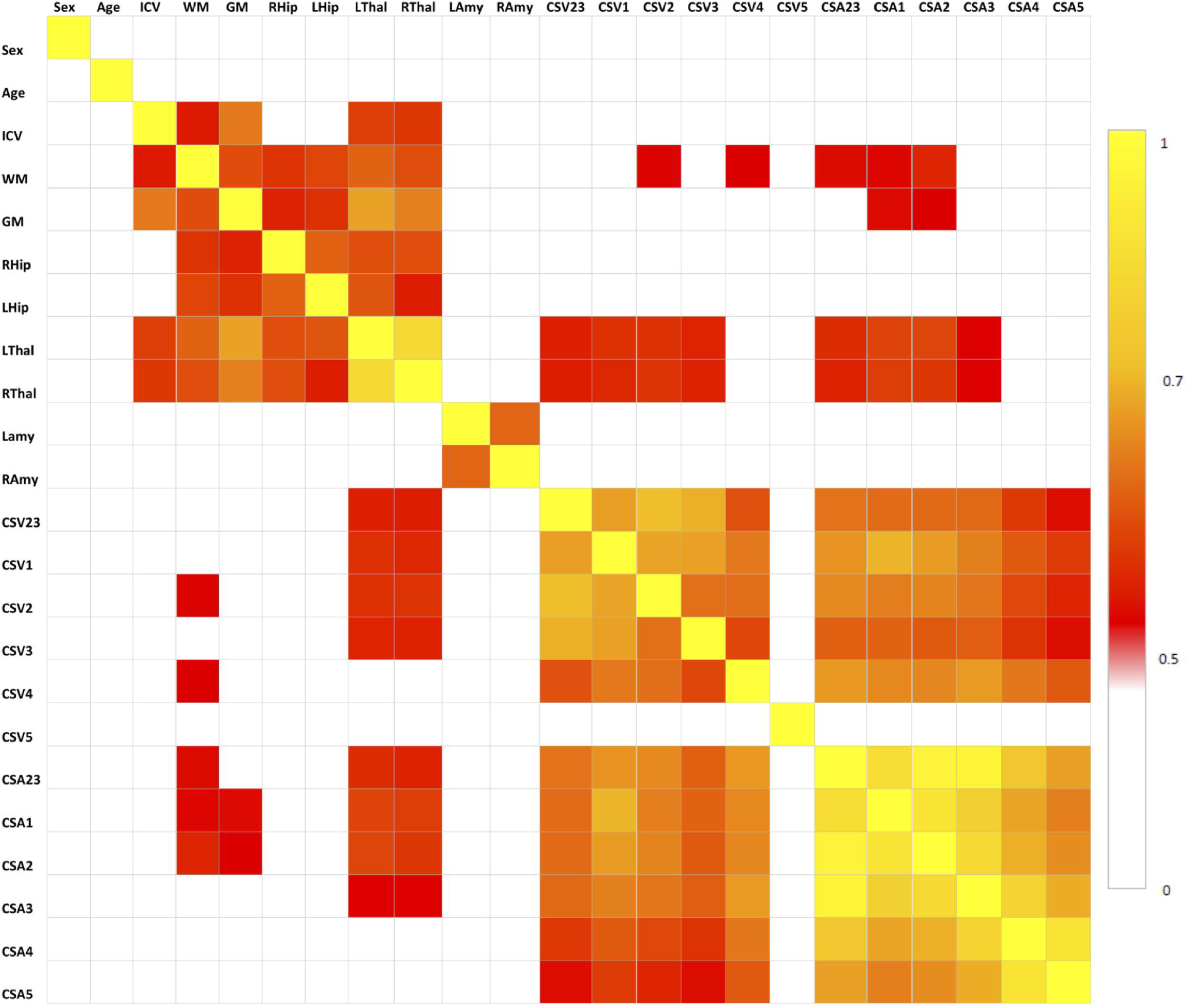
Correlation matrix between pairs of variables, tested with the Spearman’s correlation coefficient. All correlations for p<0.5 are set to white, correlations for p>0.5 are red to yellow, with yellow (p=1) being the strongest correlation. No spinal crod metrics are correlating with brain metrics with p>0.7, which is the threshold we used for extracting the set of uncorrelated features (Table 1)

Features that were considered independent from each other and that were entered in the feature selection analysis are reported in Table 4. The best features selected by the RF algorithm for the AD versus HC classification task are reported in Table 5 and include: RHip, WM, LAmy, LHip, CSA23, GM. Interestingly, CSA23 was identified as one of the most informative features to distinguish AD patients from HC. RF outcomes are reported in Table 6 and showed that the classification accuracy of AD patients is 76%, sensitivity 74% and specificity 78%. The Area Under Curve (AUC) percentage reached 86%, showing a remarkable classification performance of the RF algorithm to distinguish AD from HC subjects. Moreover, it is noticeable that the hippocampi have dominant weight, but that there is a relevant contribution to the classification from CSA23.

**Table 4:**
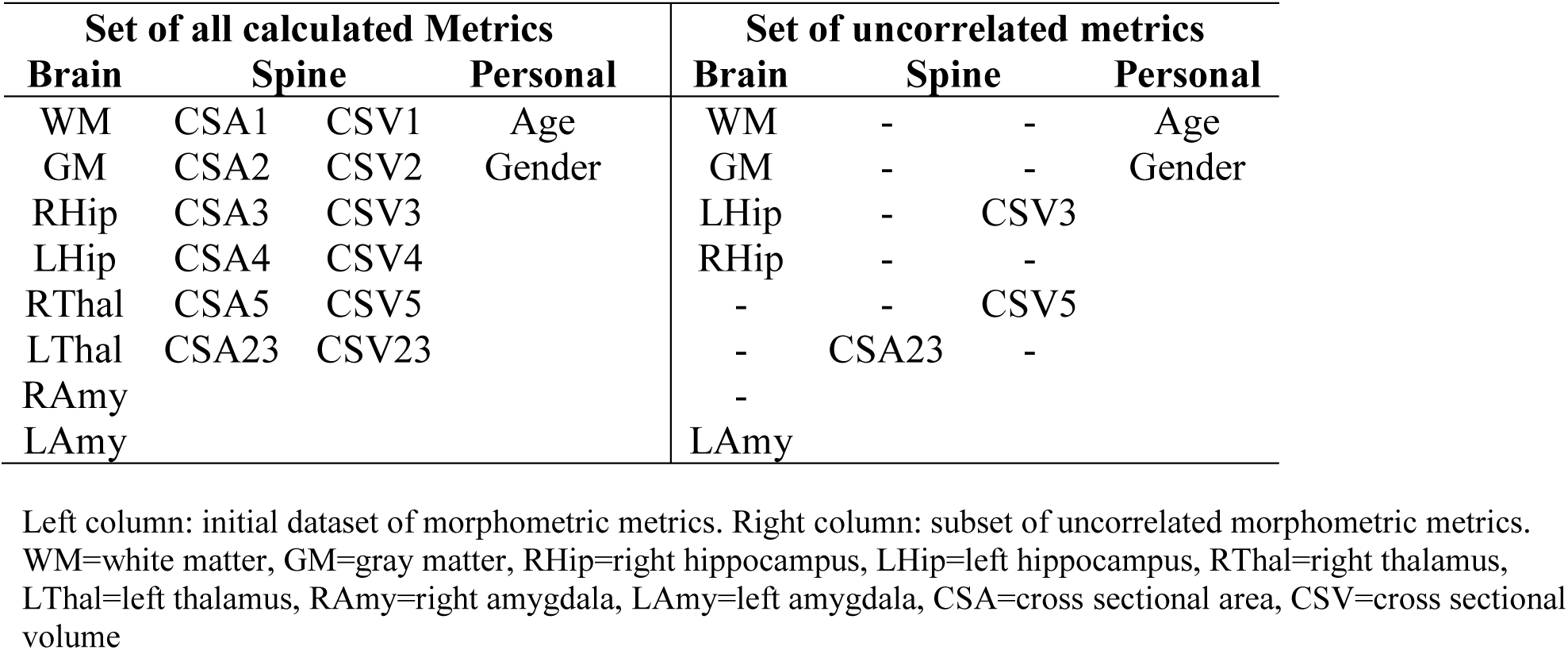
Cerebral and spinal cord morphometric metrics.

**Table 5:** Features ranking.

| Features | Weight |
| --- | --- |
| RHip | 0.1125 |
| WM | 0.0630 |
| LAmy | 0.0629 |
| LHip | 0.0615 |
| CSA23 | 0.0317 |
| GM | -0.0041 |

9 HC and 9 AD patients were used in the ranking procedure. Ranking Algorithm: ReliefF applied on a dedicated subset (30% of instances, number of neighbors = 10).

**Table 6:** Random Forest classification.

| Performance |  |
| --- | --- |
| Accuracy | 76 % |
| Sensitivity | 74% |
| Specificity | 78% |
| Area Under Curve | 86% |
| Feature | RF weight |
| LHip | 9.039 |
| RHip | 2.734 |
| LAmy | 2.263 |
| CSA23 | 1.828 |
| WM | 0.323 |
| GM | 0.060 |

23 HC and 19 AD were used to test classifier performance. A leave-one-out procedure was used to test the performance of Random Forest (RF) with the best feature subset reported in table 5. RF features weight are also reported.

### 3.4 MMSE and morphometric data relationship

The combination of the six best features, including WM, RHip, LHip, LAmy, CSA23 and GM, explained 44% of the overall variance of the MMSE. The function equation describing the linear model obtained by the regression analysis included the following terms with their weights: 0.329*LHip-0.145*RHip+0.145*LAmy+0.064*CSA23-0.227*GM+0.557*WM. The MMSE explained variance was progressively reduced by simplifying the model, i.e. removing one or more predictors, as shown in Table 7. Each separate feature significantly (p<0.005) explained a percentage of MMSE variance ranging between 13% to 36%. The feature that most explains MMSE variance was the WM volume (36%), with CSA explaining 13%.

**Table 7:** MMSE outcomes.

| Multiple Linear Model | Explained Variance | Influence Significance |
| --- | --- | --- |
| MMSE = $\beta_1$*LHip+$\beta_2$*RHip+$\beta_3$*LAmy+$\beta_4$*CSA23+$\beta_5$*GM+$\beta_6$*WM | 44% | <0.001 |
| MMSE = $\beta_1$*LHip+$\beta_2$*RHip+$\beta_3$*LAmy +$\beta_4$*GM+$\beta_5$*WM | 43% | <0.001 |
| MMSE = $\beta_1$*LHip+$\beta_2$*LAmy+$\beta_3$*GM+$\beta_4$*WM | 43% | <0.001 |
| MMSE = $\beta_1$*LHip+$\beta_2$*GM+$\beta_3$*WM | 42% | <0.001 |
| MMSE = $\beta_1$*LHip+$\beta_2$*WM | 40% | <0.001 |

| Linear Model | Explained Variance | Influence Significance |
| --- | --- | --- |
| MMSE = β*WM | 36% | <0.001 |
| MMSE = β*LHip | 30% | <0.001 |
| MMSE = β*RHip | 22% | <0.001 |
| MMSE = β*GM | 17% | 0.001 |
| MMSE = β*LAmy | 16% | 0.001 |
| MMSE = β*CSA23 | 13% | 0.005 |

MMSE Linear Regression Models. The model-explained variance is calculated with the R<sup>2</sup> index. Significance was set to p=0.05; all described model showed statistically significant influence (ANOVA).

## 4 Discussions

The present work is pioneering the investigation of spinal cord alterations in patients with dementia, and in particular with AD, a major neurodegenerative disease known for its profound effect on cognitive functions. The motor/sensorimotor system has already been shown to be affected in AD at various levels in the brain, but nobody has yet investigated the spinal cord to date^2–6^. Here we have shown that the spinal cord is very atrophic indeed, at least in established AD patients. This is an important finding, as it demonstrates that atrophy and neurodegeneration is widespread beyond areas with excellent standards such as the hippocampi and temporal lobes. Our results are, indeed, consistent with the fact that patients present significantly different brain volumes with respect to HC, and all segmented brain structures, except for the right amygdala, are statistically significantly atrophic in AD. In this context, our work goes further and demonstrates that volumes of all cervical vertebral segments are reduced in AD, with the CSV of the first and second vertebrae being significantly atrophic with respect to HC. These results are coherent with results obtained for cerebral structures and suggest the existence of a remarkable reduction (of the order of 10%) in the volume of the spinal cord in dementia. This hypothesis is further supported by significant CSA reduction for all vertebrae in patients, with CSA being calculated considering the curvature of the spinal cord^28^. Previous studies have reported spinal cord atrophy in patients with neurological diseases^40,41^, such as multiple sclerosis with focal lesions in the brain and spinal cord, but to date no studies have explored the existence of a volumetric loss of spinal cord tissue in dementia. This finding has implications for how we think about the relationship between the cognitive and sensorimotor systems, which we have shown to be conjointly affected in established AD. Not to forget that the spinal cord is also the relay of the autonomic system that has been reported as dysfunctional in AD^42–44^.

Post mortem studies of AD patients will be needed to confirm the biophysical source of spinal cord atrophy, although at first one could imagine that any change in CSA and CSV could be the result of retrograde Wallerian degeneration from the cerebral cortex^45^. Given that we have also demonstrated that spinal cord features are independent of brain volumes, it cannot be excluded that alterations in spinal cord morphometric measurements (CSA and CSV) in AD are the result of primary retrogenesis linked to myelin and axonal pathology. It is indeed very significant that a recent study of the 5xFAD animal model of AD shows amyloid plaques accumulation in the spinal cord tissue, with a particular concentration at cervical level and a time dependent accumulation that starts 11 weeks from onset; interestingly, the same study found independent and extensive myelinopathy, while the motoneurons count at 6 months was not altered compared to the wild type^46^. While we cannot be conclusive on the mechanisms of spinal cord atrophy in AD, our results are intriguing and calling for larger studies of prodromic subjects to be followed over time; such studies would also confirm whether the suggestion that the motor system (neocortex, cerebellum and spinal cord) is affected even before the cognitive one can be substantiated^3,5,14^

A further result of our work is that of all spinal cord features analysed here, the area of vertebra C2-C3 (CSA23) significantly contributes to discriminate between HC and AD patients. Usually, only atrophy of brain regions is investigated in dementia^47,48^. Indeed, spinal cord morphometric measures (CSA and CSV) alone cannot directly discriminate between AD and HC, but CSA23 was identified as one of the six best features useful to distinguish between these groups of subjects. Classifier accuracy was good and reached its best performance, around 76%, when both volume of brain structures, such as LHip and RHip (considered biomarkers of AD progression^49^), WM and GM, as well as spinal cord CSA23 were included in the classification procedure. In addition, the ROC curve between AD and HC (shown in Figure 4) reported high performance with AUC of 86%. The sensitivity and specificity of the RF algorithm, reaching 74% and 78% respectively, showed a remarkable ability in correctly identify healthy and pathological cases. Examining the RF feature weighting (reported in Table 6) it is also noticeable that CSA23 had weight higher than GM, highlighting that it should be considered as an additional biomarker together with the more conventional volumes of subcortical regions^39^. These results indicate the yet unexplored potential influence that spinal cord features can play in the classification of dementia in line with recent publications, which have recognized that other brain structures play a key role in identifying AD patients and in distinguishing between different subtypes of dementia^12,15^.

**Figure 4.**
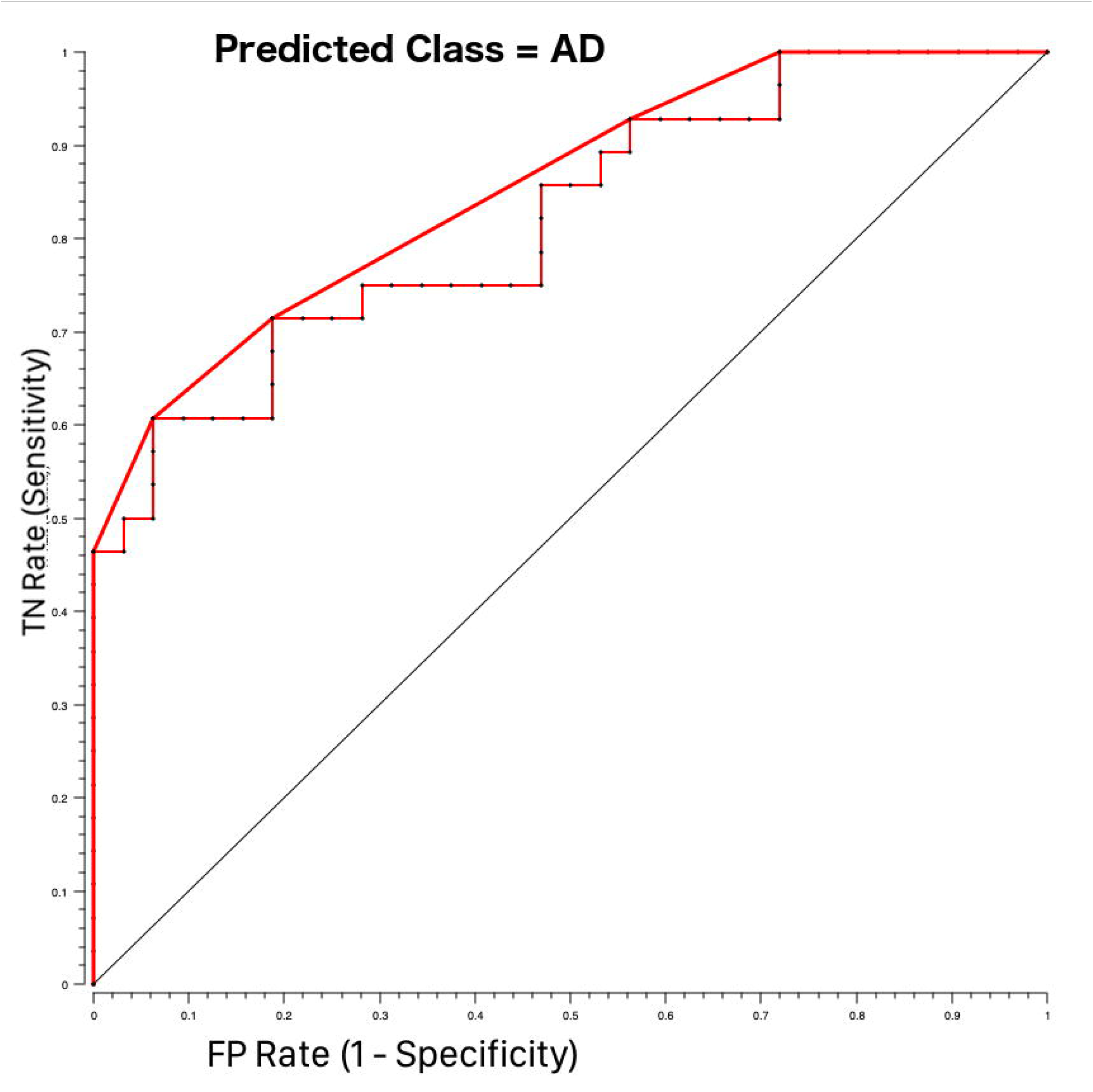
ROC curves for AD-HC classification using Random Forest feature selection. Pathological class (AD = 1) was considered as the target class. The curve shows higher performance (bold red line) than the majority algorithm (diagonal). TN rate is the rate of true negative and FP rate is the rate of false positives.

Regarding the fact that CSA23 emerged as being particularly sensitive to pathological changes in AD is in accordance with other studies in neurodegenerative diseases such as amyotrophic lateral sclerosis^17^ and could be seen as a corroborating evidence of a correlation between spinal cord atrophy and neurodegeneration. In light of the only animal model study reported to date^46^, which shows that C2-C3 is selectively affected by greater morphological biophysical alterations, our results become of remarkable value. Moreover, upper limb sensorimotor impairment is known to be clinically relevant in early AD, which is substantiating the relevance of our finding and calls for future investigations involving correlations with sensorimotor scores and purposely designed prospective studies to answer mechanistic questions.

Finally, our data show that also clinical aspects of AD are partially explained by spinal cord atrophy. Given the exploratory nature of the present study, we assessed whether spinal cord atrophy could be correlated with the variance of the MMSE, which is a global test, clinically used to assess AD severity. We found that 43% of the MMSE variance was explained with a multiple regression model implemented with all the best features included as independent variables, whereas CSA23 alone explained 13% of the MMSE variance, which is a considerable contribution indeed.

From a methodological point of view, we know that evaluating spinal cord alterations in humans in vivo is challenging due to technical and anatomical constrains, including subject positioning inside the scanner, individual subject’s neck curvature or subject’s motion. Furthermore, the spinal cord is a small structure and optimized sequences with reduced FOV and appropriate alignment should be used to obtain reproducible results^50^. Dedicated acquisition protocols would also allow one to analyse specific alterations of spinal cord GM and WM, that were not available with the present data that used 3DT1-w scans, used for whole-brain or regional brain volume calculations, to extract spinal cord features^51,52^. Regarding feature selection and classification, we know that recent studies have combined several MRI findings with machine learning approaches to attempt the classification of dementia subtypes and prediction of disease progression. Accuracy of about 80%^53,54^ was achieved when AD and HC were classified while more fluctuating results were reported when more subtypes of dementia were considered. In the present study a RF algorithm with the “leave-one-out” approach was chosen to discriminate between AD and HC because RF is robust with small numbers of subjects and performs features weighting runtime with good sensitivity and specificity.

Given the nature of this prospective study, it was not possible to investigate the involvement of the spinal cord at different stages of AD or in different types of dementia to explore its full clinical potentials. Therefore, a comprehensive battery of sensorimotor and cognitive tests should be performed to understand how the clinical and MRI pictures are evolving during the disease progression and to establish when spinal cord atrophy occurs and its clinical weight. It is also essential to promote multi-modal studies that can disentangle the contribution of myelin, amyloid accumulation, axonal swelling and axonal loss to brain and spinal cord alterations in neurodegenerative diseases to understand local and global mechanisms of damage.

In conclusion, the present work can be considered a milestone because for the first time it demonstrates in a cohort of AD and HC subjects the contribution of spinal cord atrophy to explain clinical indicators of dementia and to improve disease classification, opening also mechanistic questions for future studies. It is indeed important that we rethink in particular of how the sensorimotor and cognitive systems are affected by AD, integrating spinal cord with brain information, including the temporal lobes with the hippocampi, the motor and sensorimotor cortices, the limbic system with the amygdala and the cerebellum, which we now know are all implicated in AD.

## 7 Acknowledgements

We thank University of Pavia and Mondino Foundation (Pavia, Italy) for funding; The UK Multiple Sclerosis Society and UCL-UCLH Biomedical Research Centre for ongoing support of the Queen Square MS Centre. CGWK receives funding from ISRT, Wings for Life and the Craig H. Neilsen Foundation(the INSPIRED study), from the MS Society(#77), Wings for Life(#169111), Horizon2020(CDS-QUAMRI, #634541). This research has received funding from the European Union’s Horizon 2020 Framework Programme for Research and Innovation under the Specific Grant Agreement No.785907(Human Brain Project SGA2) for the work of FP and ED. ECTRIMS and the Multiple Sclerosis International Federation(MSIF) supported the work of GC with funding (ECTRIMS Postdoctoral Research Fellowship Program, MSIF Du Pré grant).

## 8 Authorship

CGWK, ED, FP and RL conceptualized the study. FP and RL designed and performed the analyses with support from GC. PV and NA acquired all MRI data. ES and SB acquired all neuropsychological data helping for data interpretation. AC, ES and GM enrolled all patients and performed all clinical evaluations. CGWK and ED provided support and guidance with data interpretation with clinical contribution of all physicians. CGWK, FP and RL wrote the manuscript, with comments from all other authors.

